# Can Membrane Composition Traffic Toxins? Mycolactone and Preferential Membrane Interactions

**DOI:** 10.1101/2022.05.31.494214

**Authors:** Gabriel C. A. da Hora, John D. M. Nguyen, Jessica M. J. Swanson

## Abstract

Mycolactone is a cytotoxic and immunosuppressive macrolide produced by *Mycobacterium ulcerans* and the sole causative agent of the neglected tropical skin disease Buruli ulcer. The toxin acts by invading host cells and interacting with intracellular targets to disrupt multiple fundamental cellular processes. Mycolactone’s amphiphilic nature enables strong interactions with lipophilic environments, including cellular membranes; however, the specificity of these interactions and the role of membranes in the toxin’s pathogenicity remain unknown. It is likely that preferential interactions with lipophilic carriers play a key role in the toxin’s distribution in the host, which, if understood, could provide insights to aid in the development of needed diagnostics for Buruli ulcer disease. In this work, molecular dynamics simulations were combined with enhanced free energy sampling to characterize mycolactone’s association with and permeation through models of the mammalian endoplasmic reticulum (ER) and plasma membranes (PM). We find that increased order in the PM not only leads to a different permeation mechanism compared to that in the ER membrane, but also an energetic driving force for ER localization. Increased hydration, membrane deformation, and preferential interactions with unsaturated lipid tails stabilize the toxin in the ER membrane, while disruption of lipid packing is a destabilizing force in the PM.

**STATEMENT OF SIGNIFICANCE:** Mycolactone is sole the causative agent of Buruli ulcer, a neglected tropical disease involving large necrotic lesions that can cause permanent disfigurement if left untreated. Due to its amphiphilic nature, the toxin hides from traditional diagnostic detection and the host immune system by associating with lipophilic carriers, including cellular membranes. Our work uses extensive all-atom simulations to query if the toxin has preferential interactions with different types of membranes. We find a clear preference for more disordered membranes, like the endoplasmic reticulum’s, via interactions with unsaturated lipid tails and membrane deformation. The revealed insights can be used to predict host cell distribution between different types of lipophilic carriers and to aid in the design of Buruli ulcer diagnostics.

## INTRODUCTION

Buruli ulcer (BU) disease is a debilitating skin disease caused by the bacteria *Mycobacterium ulcerans*. The disease is characterized by large necrotic lesions that surprisingly lack pain and wound healing and can lead to permanent disfigurement and disability if left untreated (1-3). While treatments for Buruli ulcer disease exist and antibiotic treatment regimens have seen striking improvements in the last few years (4, 5), early detection remains a significant challenge. This is partially because BU manifests in similar ways as other skin neglected tropical skin diseases (e.g., leprosy, leishmaniasis, and yaws) (6, 7). Another challenge is the development of potential treatments for immunosuppressed individuals, such as patients with advanced HIV coinfection or undergoing immunosuppression treatments (5). Unlike other skin neglected tropical diseases, BU is caused by a single causative agent, an exotoxin produced by the bacteria called mycolactone. This cytotoxin is secreted by the bacteria and invades host cells, where it disrupts multiple cellular processes. Its effects include impaired immune response, cell adhesion, and pain perception, as well as cell death (8). Additionally, it evades the host immune system and is difficult to detect with antibodies – making the development of diagnostics challenging (9).

Although mycolactone was initially believed to passively permeate through membranes into host cytosol (10), simulations first predicted (11) and experiments shortly thereafter demonstrated (12, 13) that it actually associates strongly with lipophilic structures such as cellular membranes, albumin, and lipoproteins. This is sensible given the toxin’s amphiphilic nature and potentially explains how being consistently carried in and transferred between lipophilic structures would effectively hide mycolactone from both innate immune processes and traditional antibody development (14). It would also explain why developed monoclonal antibodies are only effective when mixed with the toxin prior to cell exposure (14). Thus, understanding how mycolactone is distributed in and navigates the host system via lipophilic carriers would benefit the development of effective diagnostic assays, which is a noted priority of the World Health Organization (9). It would also provide fundamental insight into how bacterial amphiphiles, in general, interact with lipophilic carriers to evade the host immune system, localize, and cause disease.

Mycolactone’s effects have been shown to be mediated by its interactions with multiple cellular targets. The toxin is thought to cause analgesia through its interaction with the Angiotensin II receptor (AT2R), which is responsible for signal transduction in neurons involved in the perception of pain. The toxin binds to AT2R, triggering a signaling cascade that causes hyperpolarization of neurons which inhibits their ability to send information, thus impairing the ability to feel pain (15, 16). Skin ulceration observed in Buruli ulcer disease can be partially attributed to the toxin’s interaction with the Wiskott-Aldrich syndrome protein (WASP), a regulatory protein involved in the polymerization of actin filaments in the cytoskeleton, which enables cell adhesion to form cell tissue. The toxin binds to WASP with a high binding affinity and constitutively activates it, causing uncontrolled branching of actin filaments, defective cell adhesion, and skin ulceration (17, 18). However, most of mycolactone’s cellular effects result from targeting the Sec61 translocon (19-21), a membrane-embedded protein complex that translocates newly synthesized polypeptides into the ER for processing. The toxin binds to the protein and inhibits its translocation mechanism reducing the cell’s ability to produce many secreted and transmembrane proteins leading to immunomodulation and cytotoxic effects (8). Single amino acid mutations in Sec61 have been identified that confer human cells broad resistance to mycolactone’s cytotoxic and immunomodulatory effects while not affecting the protein’s functionality (19).

An interesting possible contributing factor to mycolactone’s impact on so many cellular processes is its strong interaction with membranes. Studies using fluorescent derivatives of mycolactone reported its passive diffusion across membranes and uptake into the cytoplasm (10). However, our previous work quantified a strong free energy of association between the toxin and model phospholipid membranes, suggesting what was previously thought of as cytosolic localization was a more likely association with the ER membrane and Sec61 translocon (11, 22). This work further demonstrated that the toxin has a strong preference for the interfacial region below the lipid headgroups, and that water molecules play a crucial role in stabilizing the polar groups of mycolactone during membrane association and permeation (22). While interacting with the membrane, mycolactone may also alter membrane properties. The toxin has been shown to perturb lipid organization in coarse-grained model membranes, suggesting it could change the formation of ordered domains (11). Experimental work using Langmuir monolayers to model the PM supports these computational predictions by demonstrating mycolactone’s interaction with the membrane at very low concentrations and the toxin’s effect on the formation of ordered microdomains (12). Collectively, these findings point to a nuanced role of membrane interactions that could be highly relevant to mycolactone’s localization and pathogenicity.

The complexity of small molecule-membrane interactions is partially due to the diversity of cellular membranes. The lipid composition of biological membranes varies at the organism, cell type, organelle, and bilayer-leaflet levels. Sphingolipids and sterols are considered to be eukaryotic lipids and differ in structure in vertebrates, plants, and fungi (23). The major phospholipid in cellular membranes, phosphatidylcholine (PC), has distinct compositions in different tissues and cell types within an organism (24). Organelles within a cell vary significantly in lipid composition and some are additionally asymmetric with different lipid composition in each leaflet. The ER membrane lacks cholesterol, has more unsaturated phospholipids, and is relatively symmetrical (25). In contrast, the PM is richer in cholesterol and sphingolipids and is asymmetric with negatively charged phosphatidylserine (PS) lipids in the cytoplasmic leaflet (25, 26). Accompanying the variations in lipid composition are different physical properties such as bilayer packing, thickness and fluidity, which influence interactions with proteins and small molecules. Given this, it is likely that mycolactone has different interactions with different types of membranes that influence its host distribution and mechanism of pathogenicity.

This work aims to characterize the interaction of mycolactone B – the cytotoxic isoform of the toxin (27) – with different model membranes to better understand membrane-specific interactions and host cell distribution. We build upon previous work, where we characterized the association of the toxin with a pure 1,2-dipalmitoyl-sn-glycero-3-phosphocholine (DPPC) membrane (22), to focus on more realistic heterogeneous model membranes representative of the PM and ER membrane. We open by describing how we use all-atom molecular dynamics (MD) to simulate the membranes to identify the existence of, and cause behind, preferential membrane association. We employed Transition-Tempered Metadynamics (TTMetaD) enhanced free energy sampling in order to calculate the permeation free energy profiles and characterize the association of the toxin with our model membranes. We then present the obtained free energy surfaces, which show significantly different permeation mechanisms in the two membranes and an association preference for the ER membrane. The PM is shown to have more ordered lipids during toxin association, which limits stabilizing polar interactions with water molecules and lipid headgroups. In contrast, more flexible lipids in the ER membrane facilitate increased hydration and membrane reorganization during toxin association and permeation, contributing to the toxin’s preferential association. Our simulations reveal how mycolactone additionally increases the disorder in saturated lipid tails, disrupting lipid packing and decreasing stabilization in the PM relative to ER membrane. We close by discussing how these trends extend past the model membranes studied herein to suggest a framework for predicting the preferential association and localization of the toxin in the complex milieu of lipophilic carries in the host environment.

## METHODS

### Transition-Tempered Metadynamics

A substantial challenge for computational studies of biological processes is that most are too slow to be ergodically sampled in available MD simulation timescales. In most systems, relevant configurations are separated by free-energy barriers that are unlikely to be crossed without the aid of adaptive or enhanced sampling techniques, such as Metadynamics (MetaD) (28, 29). In non-tempered MetaD, Gaussian bias potentials are gradually added to the system Hamiltonian to nudge the system to new regions of phase space. These biases are usually applied to a combination of simple variables called collective variables (CVs). In principle, this bias accumulates as a summation of Gaussian functions deposited along the trajectory until all regions are equally sampled (i.e., diffusive motion along the CV has been obtained). However, the combined bias will oscillate around the desired free energy profile (potential of the mean force; PMF) instead of converging to it asymptotically. The addition of untempered Gaussian functions can also destabilize the system when too much energy is added. Well-Tempered Metadynamics (WTMetaD) (30) was developed to address this issue and converge to the real free energy surface. The bias added in WTMetaD is smoothly tapered to zero with a time-dependent quantity such that the bias height decreases exponentially with respect to the local bias energy. Dama et al. (31) demonstrated that WTMetaD indeed converges the bias potential asymptotically to a linearly scaled inverse of the underlying free energy.

Nevertheless, it can be challenging to determine the optimal rate of reducing the height of the bias before the simulation. Ideally, this rate should be proportional to the free energy barrier to be crossed. This information is generally not known ahead of time. If one chooses a too rapid of a rate, the system may be trapped in local minima. Alternatively, selecting too slow of rate can destabilize the system as large biases shift sampling to high-energy irrelevant regions of phase space and cause errors in the resulting free energy profile. Transition-Tempered Metadynamics (TTMetaD) (32) was developed to avoid these regimes. It is also smoothly convergent, like WTMetaD, but does not require previous knowledge of the barrier height (32). It only requires approximate locations of the minima on either side of the barrier. The local bias energy is replaced by a global property, *V*^∗^, which is the minimal bias on the maximally biased path among all the continuous paths *s*(*λ*) connecting the preselected basins in the CV space, according to:

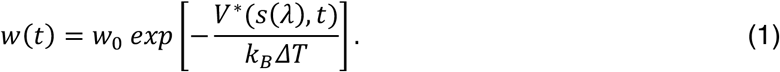

In equation 1, *w* is the height of the Gaussian (*w*_0_) divided by its deposition stride at time *t*. The rate of reducing the height of the Gaussian bias is controlled by the parameter *ΔT* (tempering). *V*^∗^ is effectively the amount of the bias energy necessary to sample the transition state (TS) region. Before the basins are connected through the TS, *V*^∗^ remains zero and the Gaussian height is not tempered. Once the basins are connected, the TS region has been traversed and the bias is aggressively tempered. Therefore, a simulation using TTMetaD efficiently fills the minima in the first step and then quickly tempers the bias to converge the free energy surface.

In our study, two-dimensional (2D) PMFs were calculated because they were more efficient in converging the desired free energy profiles and because of the increased fidelity in capturing important angular degrees of freedom previously demonstrated in membrane permeation simulations (22, 33, 34). The 2D-PMF is calculated from the reverse of the average bias energy from the independent replicas and then (diagonally) symmetrized with respect to the center of the membrane and toxin orientation (the CVs). The minimum free energy path represents the most probable pathway in the large ensemble of permeation processes, and it was calculated with a zero-temperature string method (35). Finally, the one-dimensional (1D) PMFs were obtained from the free energy along each minimum free energy path.

### Collective Variables

Identifying a set of CVs that appropriately identifies the critical slow degrees of freedom that describe the motion of complex biological processes, such as membrane permeation, can be a significant challenge. In principle, the bias energy is deposited along the CV space to increase the speed of exploration of the free energy surface. However, a poor choice of CVs can lead to slower convergence or even a complete divergence from the relevant regions of phase space. For simulations of permeation of small organic molecules through lipid membranes using TTMetaD, Sun et al. (34) have shown that 2 CVs are often necessary. The first is a *z*-component distance (*z*, CV1) between the center of the small molecule that describes the translation through the membrane, and another that describes the molecular orientation (*θ*, CV2) during the process. In our previous work on mycolactone (22), we tested the distance between the center of mass (COM) of the membrane to the COM whole molecule versus the COM of the lactone ring as possible CV1 values. The latter was better able to delineate between a wide range of configurations and demonstrated faster convergence. Thus, it was used in the current work. CV2 was defined as the angle between the vector connecting the hydroxyl groups of the end of the tails (northern and south) and the normal vector to the lipid membrane. The two CVs are represented in **Figure 1**.

**Figure 1.**
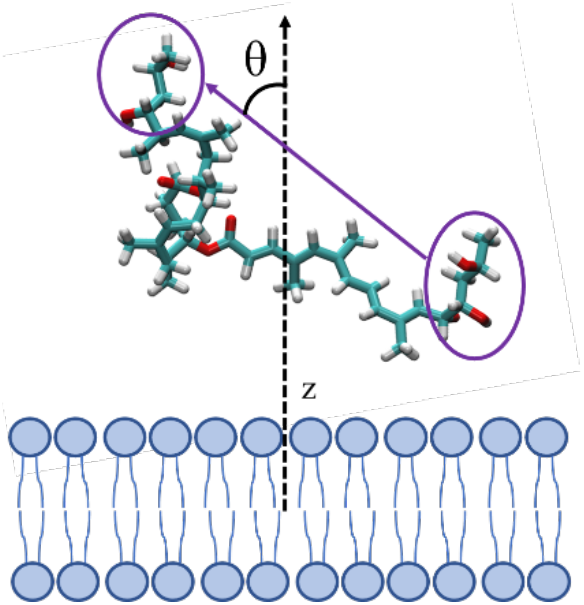
Collective variables used to define permeation where CV1 is the z-component of the distance between the COM of the membrane and that of the lactone ring, while CV2 is the angle *θ* between membrane normal and the vector connecting the hydroxyl groups of the northern (left circle) and southern chains (right circle).

### Computational Protocol

The all-atom membrane systems were built using the CHARMM-GUI Membrane Builder for Mixed Bilayers (36) with a hydration ratio of 60 for each lipid. The ER bilayer consisted of 198 units of 1-palmitoyl-2-oleoyl-sn-glycero-3-phosphocholine (POPC) and 102 units of 1-palmitoyl-2-oleoyl-sn-glycero-3-phosphoethanolamine (POPE). In addition to 80 units of POPC and 64 POPE components, the PM membrane also had 80 units of DPPC and 96 cholesterol (CHL) molecules. These numbers were chosen to represent the phospholipid compositions of the plasma and ER membranes in mammals (37) approximately. Although mammalian plasma membranes are asymmetric bilayers, our model PM is symmetric due to the challenge of preparing stable, asymmetric model membranes suitable for biophysical studies (37, 38). Likewise, existing force fields have been parametrized based on data from symmetric bilayers, thus adding a possibility of creating artifacts or significant differences in physical properties, like leaflet tension (39-42). Our PM lipid composition represents the outer cytoplasmic leaflet of mammalian plasma membranes, which is approximately two-fold more unsaturated than the inner leaflet (43). The dimensions of the simulation boxes were ∼ 9.6 × 9.6 × 9.3 and 8.4 × 8.4 × 10.4 nm for the ER and PM, respectively. For comparison to previous work (11, 22), an Amber-based force field (44) was used to describe each lipid for both bilayers with corrections to balance the hydrophobic and hydrophilic forces (45). The systems were then solvated using the TIP3P water model force field (46), heated to 310 K, and equilibrated for 250 ns until convergence was observed through property analysis (see **Figure S2**). Subsequently, one mycolactone molecule was randomly inserted into the water. For each membrane system, four replicas of mycolactone B were simulated. The toxin parameters were previously developed by Aydin et al. (22) and compared to the published results of López et al. (11). The time step to integrate the Newtonian equations of motion was 2 fs. The temperature was controlled by the canonical velocity-rescaling thermostat (47) at 310 K with a separate coupling time of 1 ps for each component (membrane – toxin and water). This temperature was chosen to mimic the human body temperature. The Berendsen barostat (48) was employed in a semi-isotropic manner (*xy*-directions coupled together while *z*-direction was independent) to maintain the pressure at 1 bar with a coupling time every 5 ps. Under periodic boundary conditions, the long-range electrostatic interactions were calculated using the smooth particle mesh Ewald method (49) with a cutoff of 1.0 nm. The short-range interaction list was updated every 10 steps with a cutoff distance of 1.0 nm. A linear constraint solver – LINCS (50) – was also applied to all hydrogen bonds. In addition to the simulated systems with enhanced sampling technique, all-atom MD simulations for both membranes systems were carried out in the absence of any biasing force to serve as a control and verify possible changes in the membrane properties.

All biased replicas were simulated for 5 µs using the GROMACS 2019.4 (51) patched with PLUMED 2.5.3 (52) to perform the enhanced method TTMetaD. The same GROMACS version was used to perform the unbiased simulations for 1 *µ*s and some of the analyses (area per lipid headgroup, membrane thickness, tail order parameter, center-of-mass distances). In order to investigate possible finite-size effects, we also performed unbiased simulations with larger membranes, doubling the number of lipids for each membrane. *Python* and *Tcl* in-house developed scripts were used to do the 2D and 1D-PMFs and hydration analysis. MDAnalysis (53, 54) was used to read the trajectories and conduct contact analysis. Visualization of the simulation boxes and trajectories and generation of the figures were done with VMD (55).

## RESULTS AND DISCUSSION

### Toxin Permeation

Two-dimensional PMFs of the permeation process with each membrane obtained from TTMetaD simulations are shown in **Figure 2**. The minimum free energy path, traced by a black line, shows the most probable permeation pathway, while inserted snapshots show dominant configurations in a few key locations. Energy minima in the PMFs correspond to configurations where the polar hydroxyl groups at the ends of the mycolactone tails are hydrogen bonding with oxygen atoms in the lipid headgroups or glycerol. At the same time, the macrolide ring is buried in the hydrophobic region, interacting with lipid tails and cholesterol molecules. These observations are consistent with our previous computational work (22), where we characterized the interaction of mycolactone with a pure DPPC model membrane.

**Figure 2.**
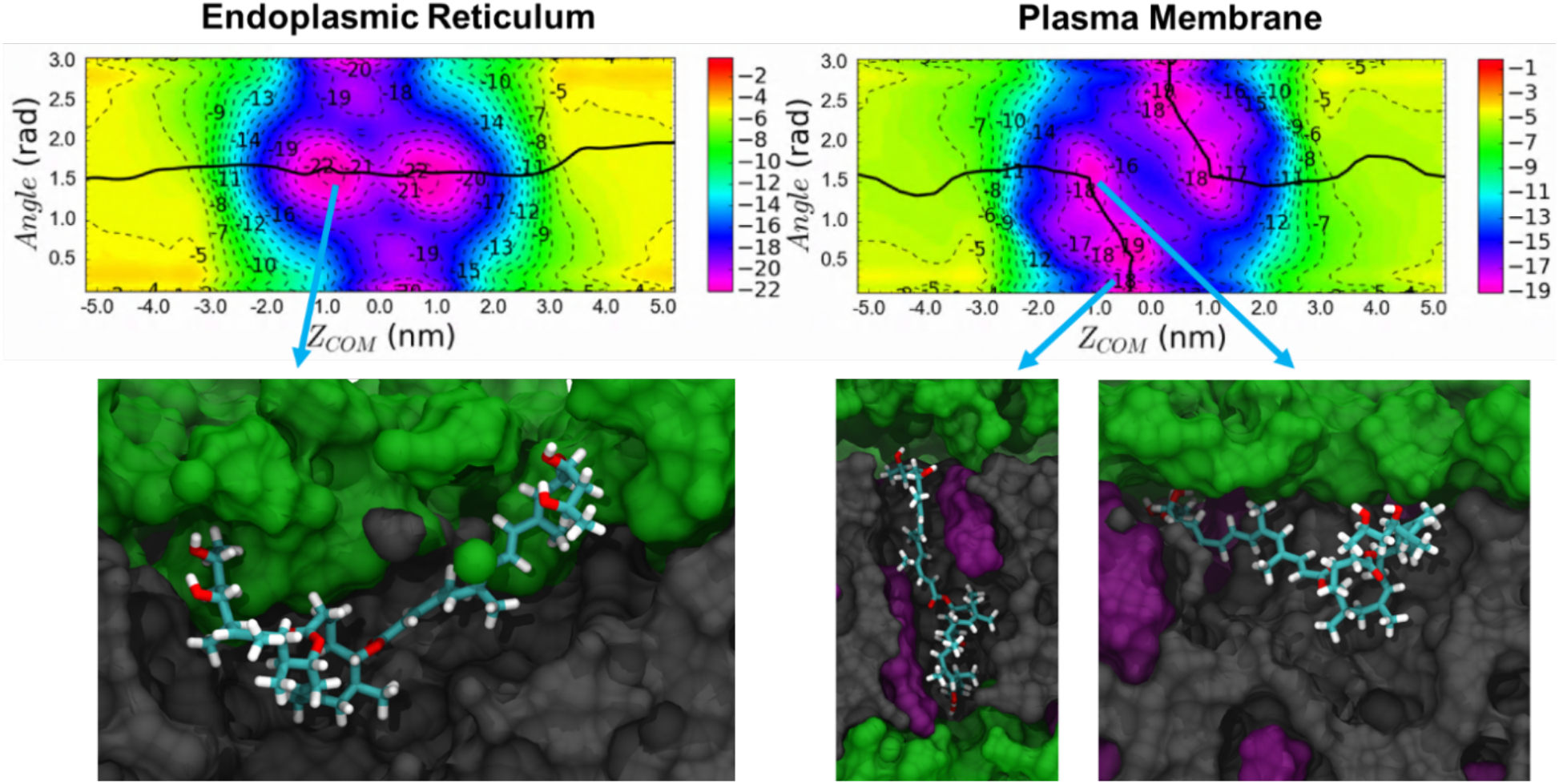
2D permeation free energy profiles through ER and PM. The minimum free energy paths (black lines) track the most common permeation pathways. Bottom figures show representative configurations in the most stable regions. The toxin is colored by atom type, while lipid headgroups, tails, and cholesterol are colored green, gray, and purple, respectively. The membranes approximately span -2.0 < Z < 2.0 nm. The free energy is shown in kilocalories per mole.

However, the minimum free energy paths also show a significant difference in the dominant permeation mechanism of the toxin through the two membranes. In the ER membrane, the toxin adopts a curved conformation at the interface below the lipid headgroups and permeates from one leaflet to the other by swapping the polar interactions of both of its hydrophilic tails from the lipid headgroups of one leaflet over to the opposite leaflet. This permeation pathway is facilitated by the deformation of the bilayer due to strong polar interactions between the toxin and lipid headgroups or water molecules (the role of hydration is further discussed below). **Figure 3** depicts the inward bend of the ER membrane allowing the toxin to continue its interactions with lipid headgroups and water molecules even at the center of the membrane. This magnitude of deformation and hydration is present in ∼30% of the conformations in this region; others have similar but fewer deep interactions with headgroups and/or water. In contrast, the permeation pathway observed in the PM has a higher and wider energy barrier in the center of the bilayer for curved conformations (0 nm, ∼1.5 radians in **Figure 2**). Thus the toxin takes a significantly different path as it permeates, swapping the interaction of its tails with the lipid headgroups one at a time and adopting an extended configuration where it interacts with both leaflet headgroups before fully migrating to the opposite leaflet. To validate the free energy landscapes of the two-dimensional PMFs, the distributions of the CVs were analyzed in unbiased simulations of each membrane system. They were found to be consistent with the energy minima of the PMF of both systems (**Figure S1**). Even in unbiased simulations of the PM, the toxin transitions between the curved and extended conformations, whereas, in the ER membrane, it remains in a curved conformation – consistent with the 2D PMFs.

**Figure 3.**
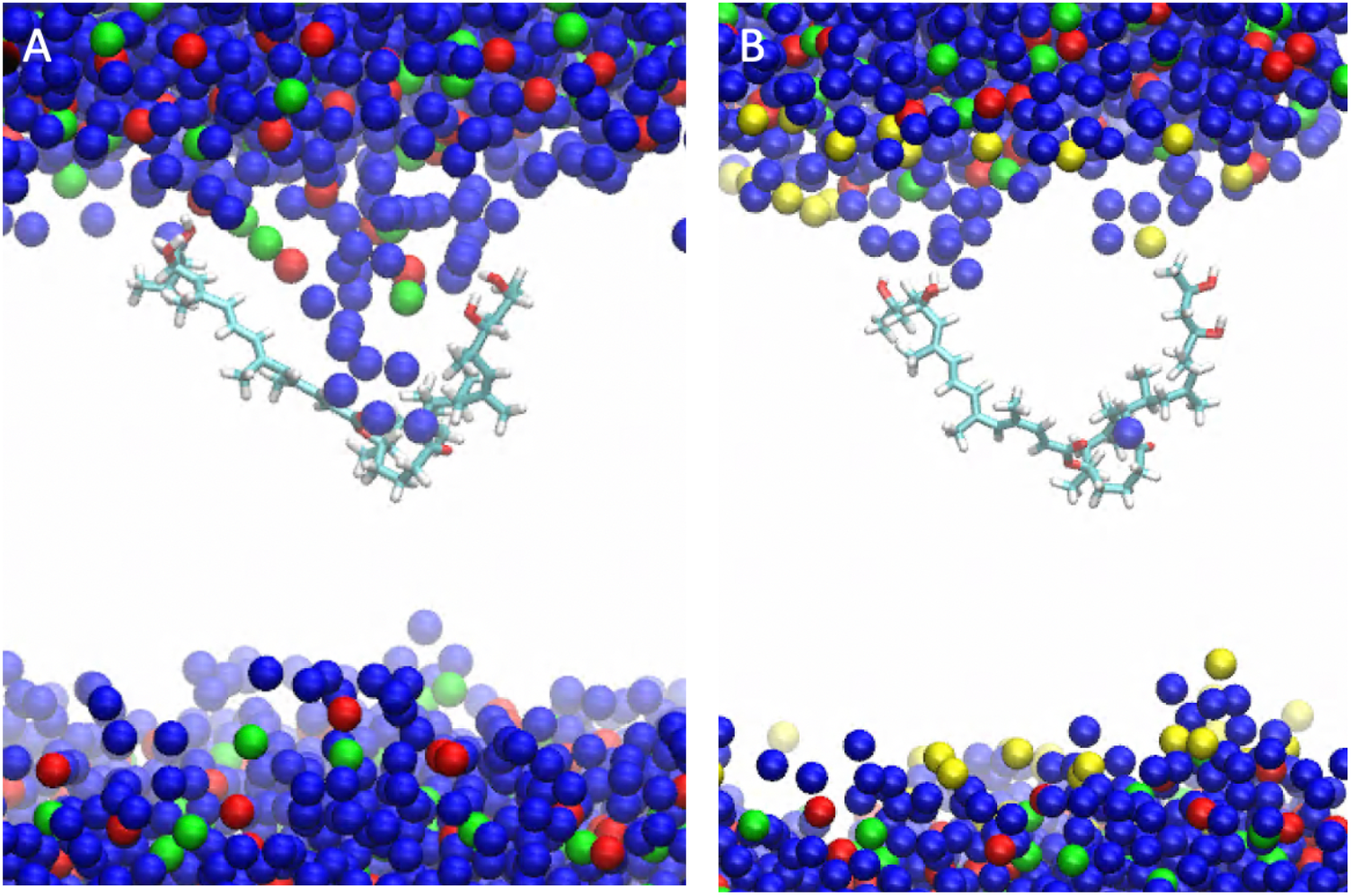
Representative configurations at Z = 0 nm and Angle = 1.5 radians (Figure 2) showing how mycolactone causes more membrane curvature and hydration defects in the ER membrane (A) compared to the PM (B). Representations include toxin (atom type), water (blue), lipid headgroup phosphorus (red), lipid headgroup nitrogen (green), and cholesterol oxygen (yellow). Lipid tails are omitted for clarity.

The difference in permeation mechanisms is likely due to the presence of cholesterol in the PM, which increases bilayer thickness and degree of order in the lipid tail region. Increased order in the PM limits the membrane deformation that stabilizes polar interactions in the pathway observed in the ER membrane (**Figure 2**). An increased bilayer thickness would additionally demand a larger degree of deformation for the toxin to retain polar interactions through the membrane center. Lipid tail order parameters and bilayer thickness of the model membranes were analyzed to verify the effect of cholesterol on membrane properties. The tail order parameter (*S*_*CD*_) of each phospholipid is a measure of the ordering and orientation of the phospholipid tail with respect to the bilayer normal, and was calculated using:

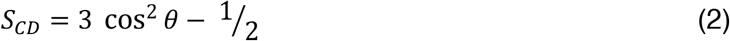

Where the *θ* is the angle between the vector of the bilayer normal and the C-H bond in the lipid. The membrane thickness was calculated as the average distance between the COM of the phosphates of the lipid headgroups of each leaflet. The results indicate that the PM is indeed thicker than the ER membrane (46.1±1.3 Å vs. 39.2±1.3 Å, respectively, where the ± indicates standard deviation) and more ordered (**Figures S2 & S3**). The influence of cholesterol on membrane order and thickness, as well as an increased height and width of a permeation free energy barrier, are consistent with previous membrane permeation studies (56-58).

### Toxin-Membrane Association and Hydration

In addition to a difference in the permeation mechanisms, the binding affinity of the toxin for the ER membrane is more favorable than it is for the PM. This is more obvious in the 1D PMFs shown in **Figure 4**. While it is helpful to quantify this difference, a more relevant question is why—*why is the toxin more stable in the ER membrane?* The preference seems to be partially due to increased interactions with water in the ER membrane relative to the PM, minimizing unfavorable interactions between the hydrophobic lipid tails and polar moieties of the toxin. Water coordination was shown in our previous work (22) to play an important role in the association of mycolactone with lipid bilayers, satisfying polar interactions as the toxin migrates into hydrophobic tail regions. Water coordination was analyzed by calculating the coordination number as a function of the z-component distance between the lactone ring and membrane center (**Figure 5**). A contact is defined as any COM distance between an oxygen atom of water and any oxygen atom of mycolactone less than or equal to 3.04 angstroms. Similar to results in our previous work (22), the toxin can interact with water molecules throughout permeation, even when the toxin is near the center of both membranes. However, mycolactone shows a higher probability of coordinating with more water molecules in the ER membrane than in the PM. This is further supported by a stronger average interaction energy between the toxin and water molecules in the ER vs. PM (−31.54 and -19.66 kcal/mol, respectively, as shown in **Table 1**). *gmx energy* was used to calculate these from the unbiased simulations using all configurations in the dominant minima in the free energy landscape (**Figure 2 & S1**). Increased hydration of the ER membrane-associated toxin can also be attributed to the presence of cholesterol and more ordered lipids in the PM, rigidifying the bilayer and condensing the lipid tails making it more difficult for the toxin to facilitate water penetration into the membrane. Other relevant interaction energies (due to toxin-toxin intramolecular interactions and toxin-lipid interactions) are stronger in the PM (**Table 1**); however, the sum of all the energies points to a stronger affinity for the ER membrane.

**Figure 4.**
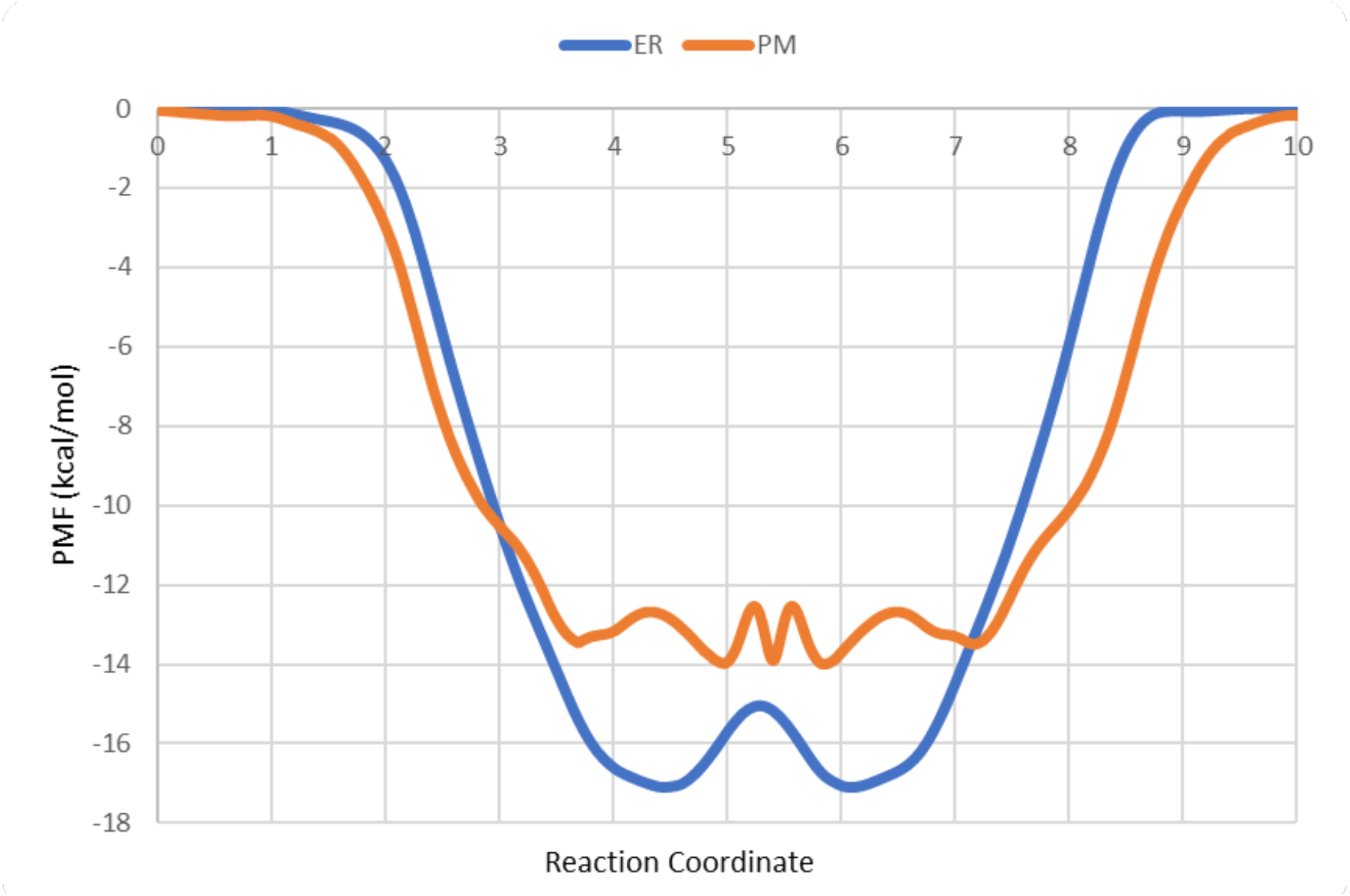
1D free energy profiles of each system obtained by plotting the minimum free energy path. The standard errors range from 0.06 to 1.22 kcal mol^-1^ for ER and from 0.02 to 2.18 kcal mol^-1^ for PM. Error bars are not shown for clarity.

**Figure 5.**
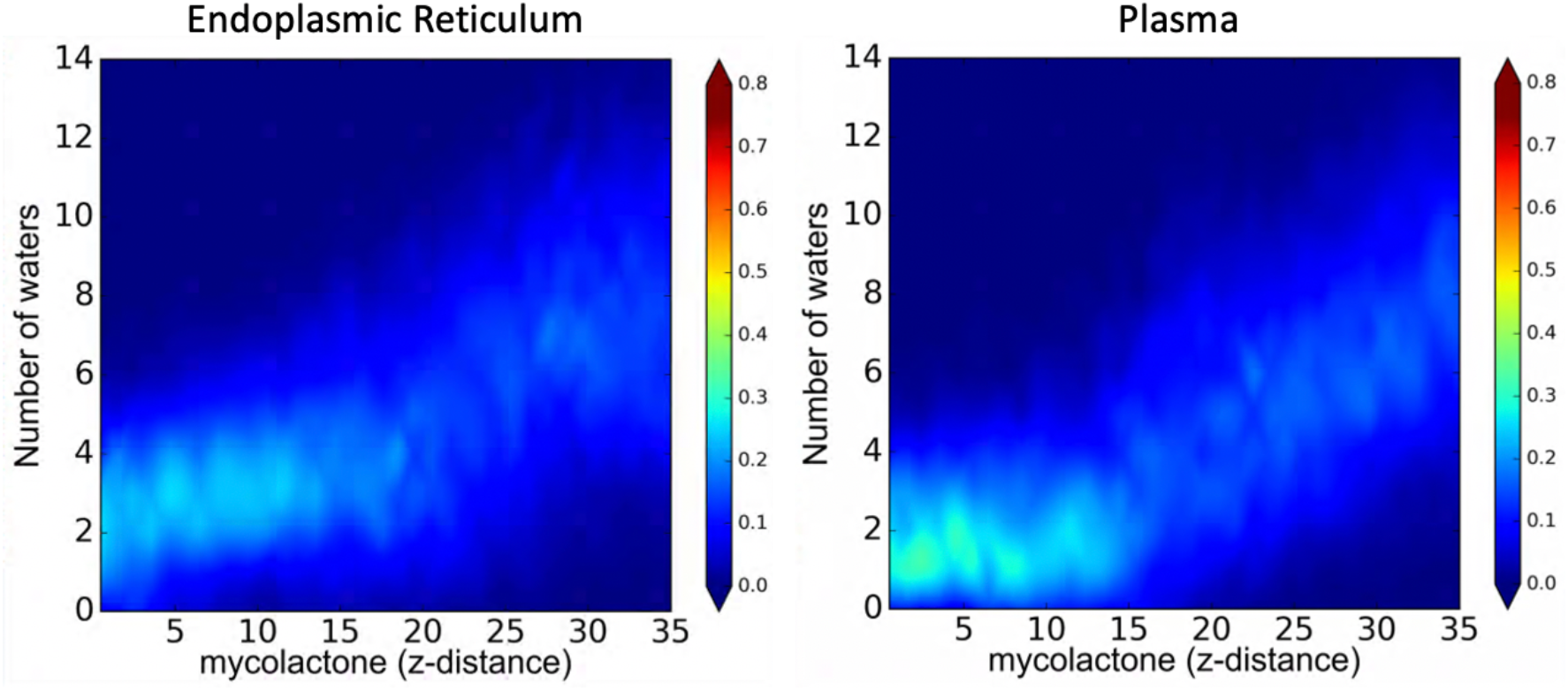
Probability distribution of the number of water molecules interacting with mycolactone with respect to the toxin position in each membrane for the biased simulations. Z-distance is shown in angstroms.

**Table 1.**
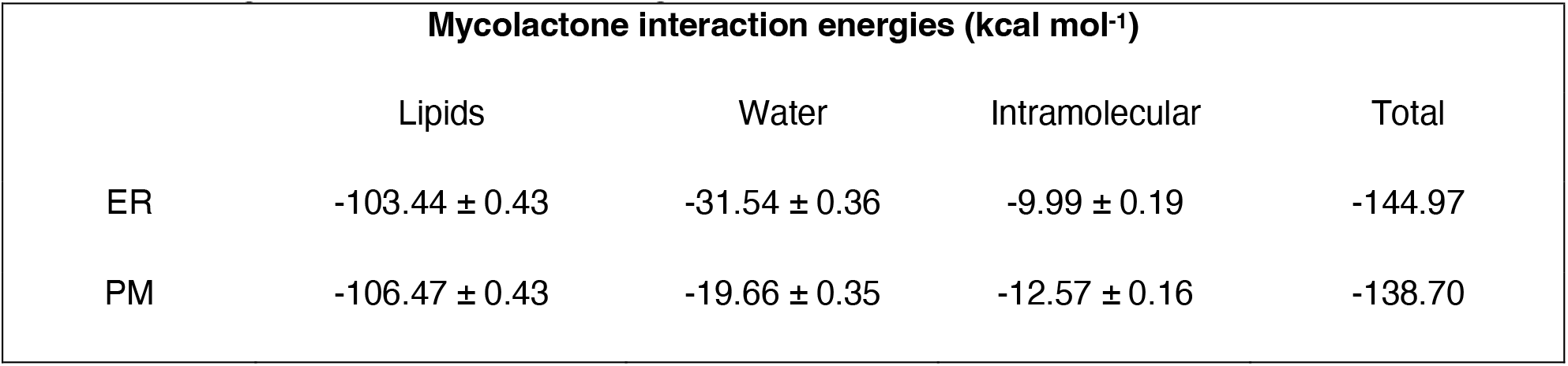
Average interaction potential energy of mycolactone with different components of the system.

| Mycolactone interaction energies (kcal mol<sup>-1</sup>) |  |  |  |  |
| --- | --- | --- | --- | --- |
|  | Lipids | Water | Intramolecular | Total |
| ER | -103.44 ± 0.43 | -31.54 ± 0.36 | -9.99 ± 0.19 | -144.97 |
| PM | -106.47 ± 0.43 | -19.66 ± 0.35 | -12.57 ± 0.16 | -138.70 |

Another contributing factor to the toxin’s preference for the ER membrane is the entropic cost of restricting the toxin’s conformational freedom in the more rigid PM. The more compact PM expectedly reduces the conformational freedom of the toxin in the bilayer. To estimate this effect, GROMACS tools (*gmx covar* and *gmx anaeig*) were used to estimate the configurational entropy of mycolactone based on Schlitter’s formula (59) and reported higher configurational entropy of the toxin in the ER membrane than in the PM (137.07 kcal mol^-1^ vs. 134.36 kcal mol^-1^, respectively).

### Lipid-Lipid Interactions

To further investigate mycolactone’s effect on the physical properties of the ER and PM, tail order parameters were calculated for phospholipids near the toxin (within 5 angstroms) in unbiased simulations and compared to tail order parameters of membrane simulations without mycolactone (**Figure S4**). In both membranes, the toxin disrupts the order of local lipids, but the effect is more significant in the PM. The larger decrease in tail order parameters near mycolactone in the PM suggests the toxin disrupts lipid packing more substantially in this membrane.

To see if this disruption has an additional ER-favoring enthalpic cost due to lost lipid-lipid interactions, the average enthalpic interaction energies for lipids were computed using *gmx energy*. Average lipid-lipid interaction energies were calculated from unbiased simulations of the ER and PM both with and without mycolactone (**Table 2**). In the system with mycolactone, the average interaction energy was calculated for lipids near the toxin (within a cutoff of 20 Å) and far from the toxin (above a cutoff of 40 Å). The difference in lipid-lipid interaction energies between the lipids near and far from toxin was indeed found to be greater in the PM. However, when comparing to the membrane-only systems, it’s apparent that the local disruption for the ER and PM are equivalent (52.3 +/- 0.5 kcal/mol). The near/far difference actually comes from a greater increase in enthalpic interactions for the lipids far from the toxin in the PM. This was at first counterintuitive and encouraged us to dig deeper to understand the origin of the near/far difference.

**Table 2.** Average interaction potential energy between lipids of the ER membrane and PM in systems with the toxin and systems without*.

| Lipid-Lipid interaction energy (kcal mol<sup>-1</sup>) |  |  |  |  |  |
| --- | --- | --- | --- | --- | --- |
| | Near the toxin | Far from the toxin | $\Delta_{(\text{near-far})}$ | Membrane without toxin | $\Delta_{(\text{near-without})}$ |
| ER | -144.91 ± 0.69 | -209.16 ± 0.45 | 64.25 | -196.63 ± 0.04 | 51.72 |
| PM | -89.94 ± 0.14 | -169.37 ± 0.24 | 79.43 | -143.08 ± 0.10 | 53.14 |

In order to verify that these observations were not influenced by the size of our simulations, larger unbiased simulations were run for both the ER and PM systems. The systems were created by doubling the number of lipids and treated with the same computation protocol (see Methods). The analyses (**Table S1, S2**, and **Figures S2, S6**) of these systems are consistent with the results described above and below, suggesting size artifacts are not significantly altering the reported properties.

We next probed whether or not mycolactone has preferential interactions with the specific lipid components making up our model membranes. The lateral radial distribution function (RDF) of mycolactone was computed with respect to each lipid component in unbiased simulations. The RDFs from the ER membrane (**Figure S5**) show a slight increase in the local density of POPC and a decrease in POPE lipids around the toxin. Given that these lipids share the same tails (one saturated and one monounsaturated), POPC is likely locally increased by the cost of packing defects. With the larger choline group, POPC would more effectively minimize the packing defects introduced by the toxin.

Interestingly, the PM RDFs (**Figure S5, S7**) show that the toxin pulls the same lipids (POPC and POPE) into its local interaction zone and pushes the saturated DPPC as well as cholesterol out. Although POPC’s role in decreasing packing defects still contributes to this distribution, the main driving force is likely the toxin’s preference for unsaturated lipids, which is consistent with findings in a previous computational study (11). In that study, coarse-grained molecular dynamics simulations were used to study the interaction of mycolactone with pure and mixed model membranes. The results suggested that in a membrane consisting of a ternary mixture of saturated lipids, unsaturated lipids, and cholesterol, the toxin preferentially interacts with unsaturated lipids in a liquid-disordered domain over-saturated lipids and cholesterol in a liquid-ordered domain.

Thus, mycolactone disrupts local lipid-lipid enthalpic interactions equivalently in the PM and ER because it induces a very similar local composition. This, in turn, increases the packing of saturated lipids just outside of the local toxin environment in the PM, making the PM association more enthalpically favorable in terms of lipid-lipid, lipid-toxin and toxin intramolecular interactions (**Tables 1 & 2**). However, the entropic cost of ordering the membrane would counteract this. Collectively, the energetic driving forces biasing the toxin to associate with ER over the PM are largely entropic, with the addition of more favorable water-toxin enthalpic interaction energies in the ER membrane.

It is important to consider that mycolactone’s preference for the unsaturated lipids of our model PM also suggests that the toxin would readily transition from the outer to the inner leaflet of mammalian PMs, which is less ordered and more unsaturated relative to the outer leaflet (43). A localization to the inner leaflet of the PM might facilitate transitions from the PM to the ER membrane via plasma-ER membrane contact sites (60) or to WASP, which is tethered to the inner leaflet of the PM.

## CONCLUSIONS

This work focused on mycolactone, an amphiphilic exotoxin that causes the pathogenesis of Buruli ulcer disease by invading host cells and binding to intracellular targets to induce cell death. Our motivation was to better understand the toxin’s localization in the host to aid in the development of effective diagnostics and provide insight into the toxin’s pathogenesis. Previous experimental and computational work has reported a strong association affinity between mycolactone and lipid bilayers (11, 12, 22). The specificity of this interaction, however, was unclear. In this work, we characterized the interaction of mycolactone with heterogeneous model plasma and ER membranes. Given the size of mycolactone and the depth of the association minima, we combined all-atom molecular dynamics simulations with TTMetaD enhanced free energy sampling in order to fully converge permeation free energy profiles. The resulting free energy profiles capture the anticipated strong association with both membranes, where dominant minima consist of configurations that satisfy polar and non-polar interactions of the toxin well under the lipid headgroups.

Our simulations demonstrate that lipid composition influences both membrane association and permeation for mycolactone. The dominant permeation pathway observed in the PM involves an extension of the toxin’s two polar tails across the bilayer, while that in the ER membrane retains a curved c-shape conformation. This difference is due to the presence of cholesterol in the PM, which in turn causes increased rigidity and bilayer thickness. Thus, it is more difficult for the toxin to deform the PM and retain polar interactions with lipid headgroups and water molecules during permeation through the membrane mid-plane. The free energy profiles also indicate a stronger affinity of mycolactone for the ER membrane, which can again be traced to the presence of saturated lipids and cholesterol in the PM. Increased order and lipid packing limit hydration and interactions with polar headgroups. The influence of hydration is significant, leading to an enthalpic bias towards the ER due to toxin-water interactions that are actually stronger than the toxin-lipid and toxin-toxin intramolecular interaction energies, both of which favor the PM. Additionally, the more rigid PM restricts the toxin’s conformational flexibility – making association with the ER membrane more entropically favorable. Furthermore, mycolactone disrupts lipid-lipid interactions in the PM more than in the ER, drawing the unsaturated lipid components (POPC and POPE) into its local environment and pushing the saturated lipid (DPPC) and cholesterol out. Thus, diagnostic design principles should target disordered lipophilic carriers with unsaturated lipid components.

The predicted stronger affinity for the ER membrane is consistent with mycolactone’s rapid accumulation in the ER following cellular uptake and its primary targeting of the ER membrane-embedded Sec61 translocon. However, it is unlikely that equilibration alone explains the toxin’s trafficking to the ER membrane. Rather some form of active uptake mechanism via endocytosis or lipid transfer proteins may also be at play. Moreover, the more considerable disruption of lipid-lipid interactions in the PM combined with the toxin’s preference for unsaturated lipids might suggest that mycolactone has a predilection for the PM inner leaflet and for liquid-disordered regions over liquid-ordered regions of biological membranes that exhibit lateral heterogeneity, which could play a role in the toxin’s trafficking to WASP near regions of actin branching.

Collectively, these findings provide a better understanding of mycolactone’s lipid-specific interactions and localization, which could facilitate the design of ideal lipophilic ‘sponges’ with high toxin affinity as diagnostic or laboratory tools for Buruli ulcer disease. More generally, increasing our understanding of mycolactone may provide clues into how other pathogenic amphiphiles, a class of pathogen associated molecular patterns that are more difficult to target, navigate host systems via lipophilic carriers. Future work should focus on asymmetric model membranes, how membrane curvature influences association, the interaction of the toxin with other lipophilic carriers such as lipoproteins and albumin, and potential active uptake mechanisms. In addition to theoretical modeling, experimental analyses of the toxin’s interactions with different lipophilic carriers, potentially with techniques using asymmetric giant unilamellar vesicles (aGUVs) (61), will be of great value.

## SUPPORTING MATERIAL

Supporting Material can be found online at https://doi.org/XX.YYYY/j.bpj.2022.0Z.ZZZ

## AUTHOR CONTRIBUTIONS

J.M.J.S., G.C.A.H and J.D.M.N designed the research. G.C.A.H and J.D.M.N performed the simulations and analyses. All authors interpreted the results and wrote the manuscript.

## ACKNOWLEDGEMENTS

The authors thank Professor Rich Pastor and Jeffrey Klauda for helpful discussions. We gratefully acknowledge support from the National Institute of General Medicine of the National Institutes of Health under award number R35GM143117 and computational support from the Extreme Science and Engineering Discovery Environment supported by the National Science Foundation (Grant No. ACI-1548562) under allocation MCB200018 as well as the Center for High Performance Computing at the University of Utah.

