## Supporting Material for "Can Membrane Composition Traffic Toxins? Mycolactone and Preferential Membrane Interactions"

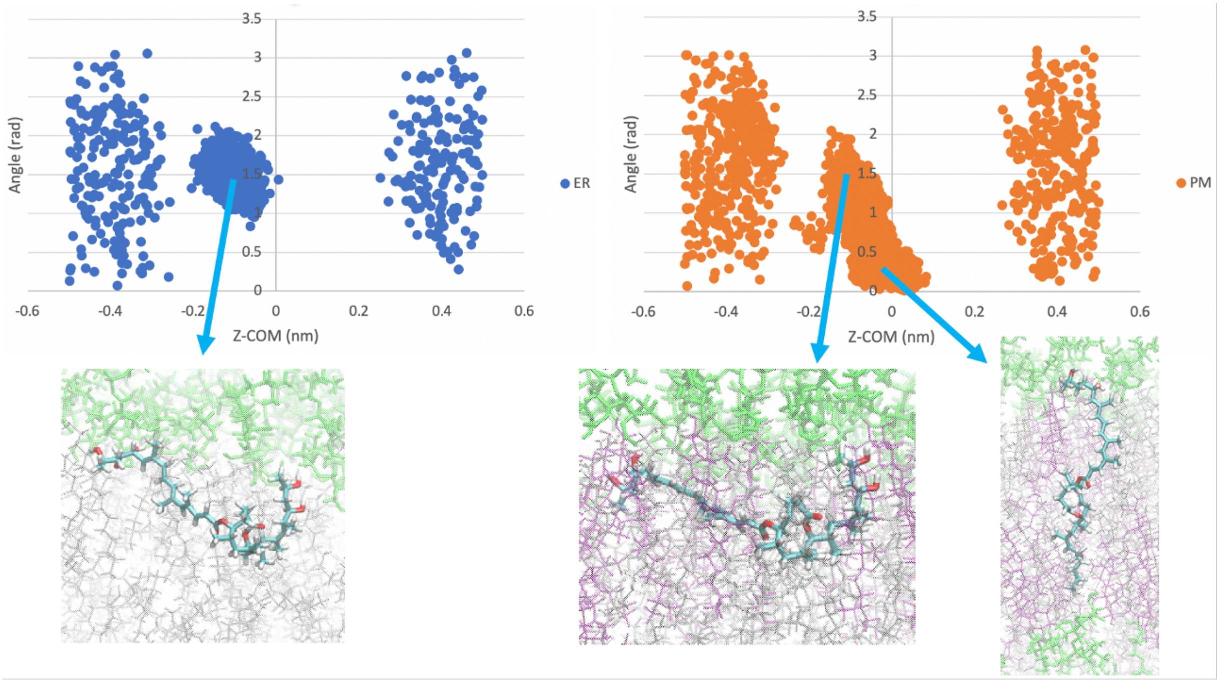


**Figure S1**. Distribution of the two CVs in unbiased simulations of mycolactone with the ER and plasma membranes (PM). Snapshots highlight the dominant configurations found in the most probable regions.

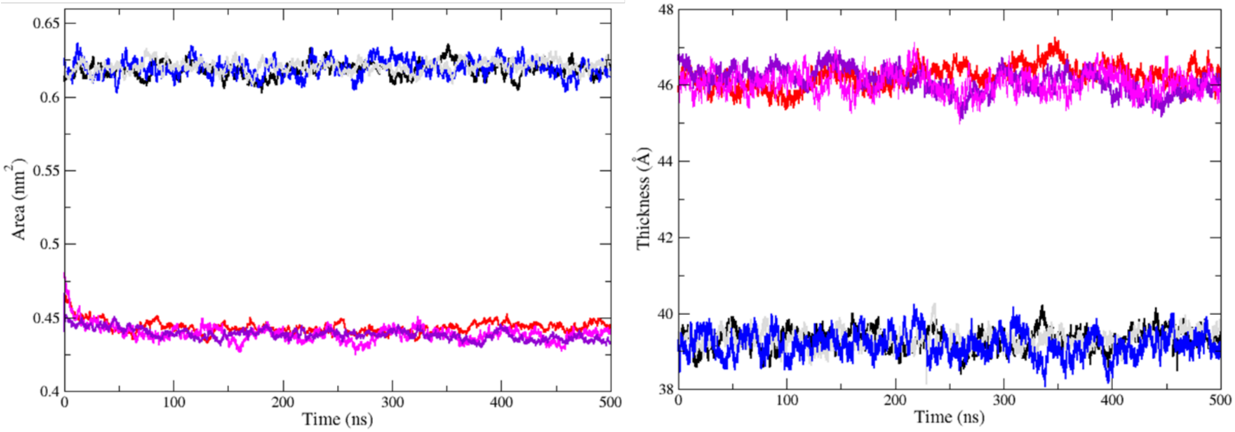


**Figure S2** Area per lipid (APL) head group (left) and membrane thickness (right) of the unbiased toxin plus membrane and pure membrane systems. APL is calculated by dividing the x,y area of the simulation box by the number of lipids in a leaflet. Mixed membranes bilayers don't have experimental reference values. Membrane thickness is calculated by the distance between the COM of phosphorous atoms in the headgroups of each bilayer leaflet. Colors: large ER in black, large ER + toxin in gray, smaller ER in blue, large PM in red, large PM + toxin in violet, smaller PM in magenta.

#### SUPPORTING MATERIAL

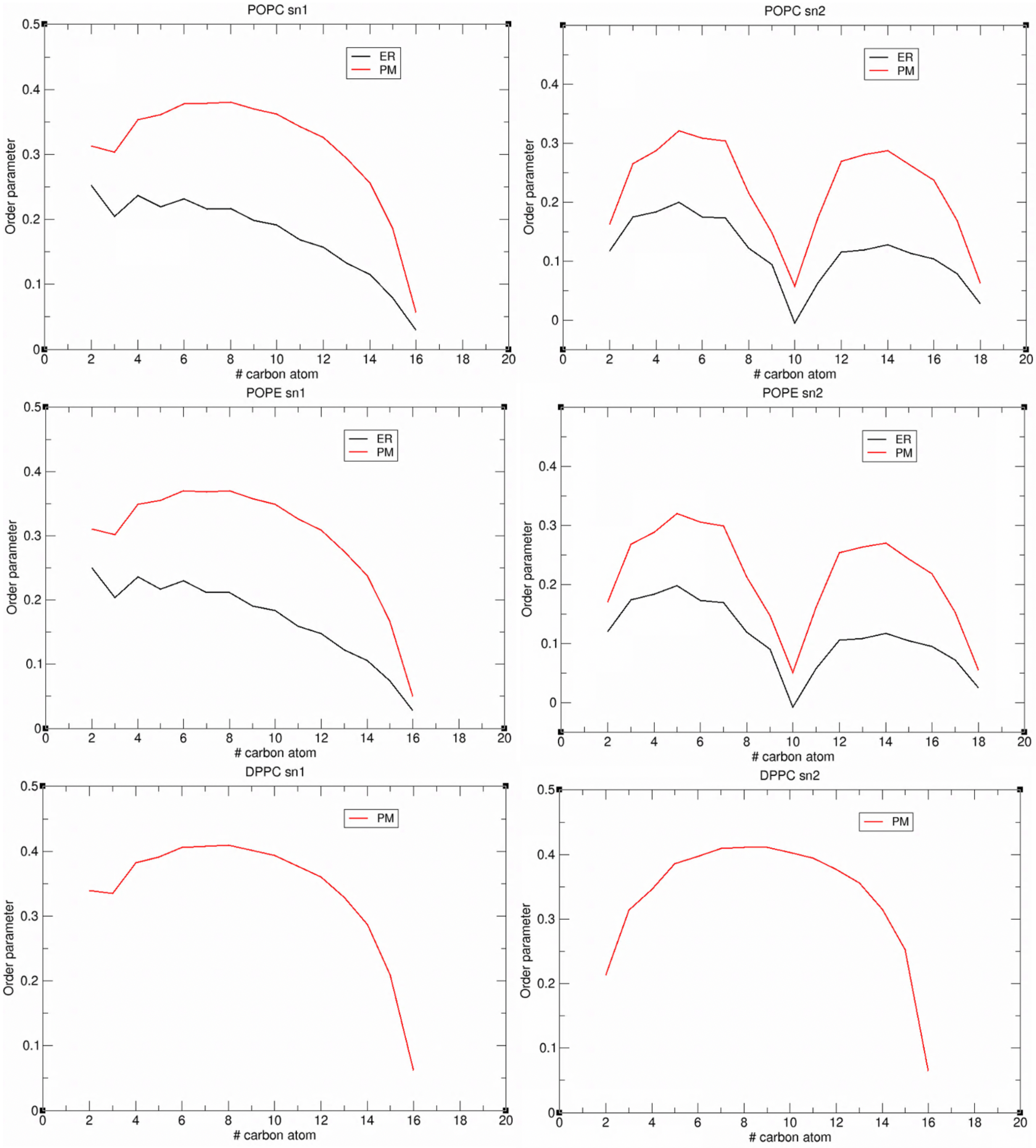


**Figure S3**. Tail order parameters of each phospholipid making up the ER and PM in membrane-only simulations.

#### SUPPORTING MATERIAL

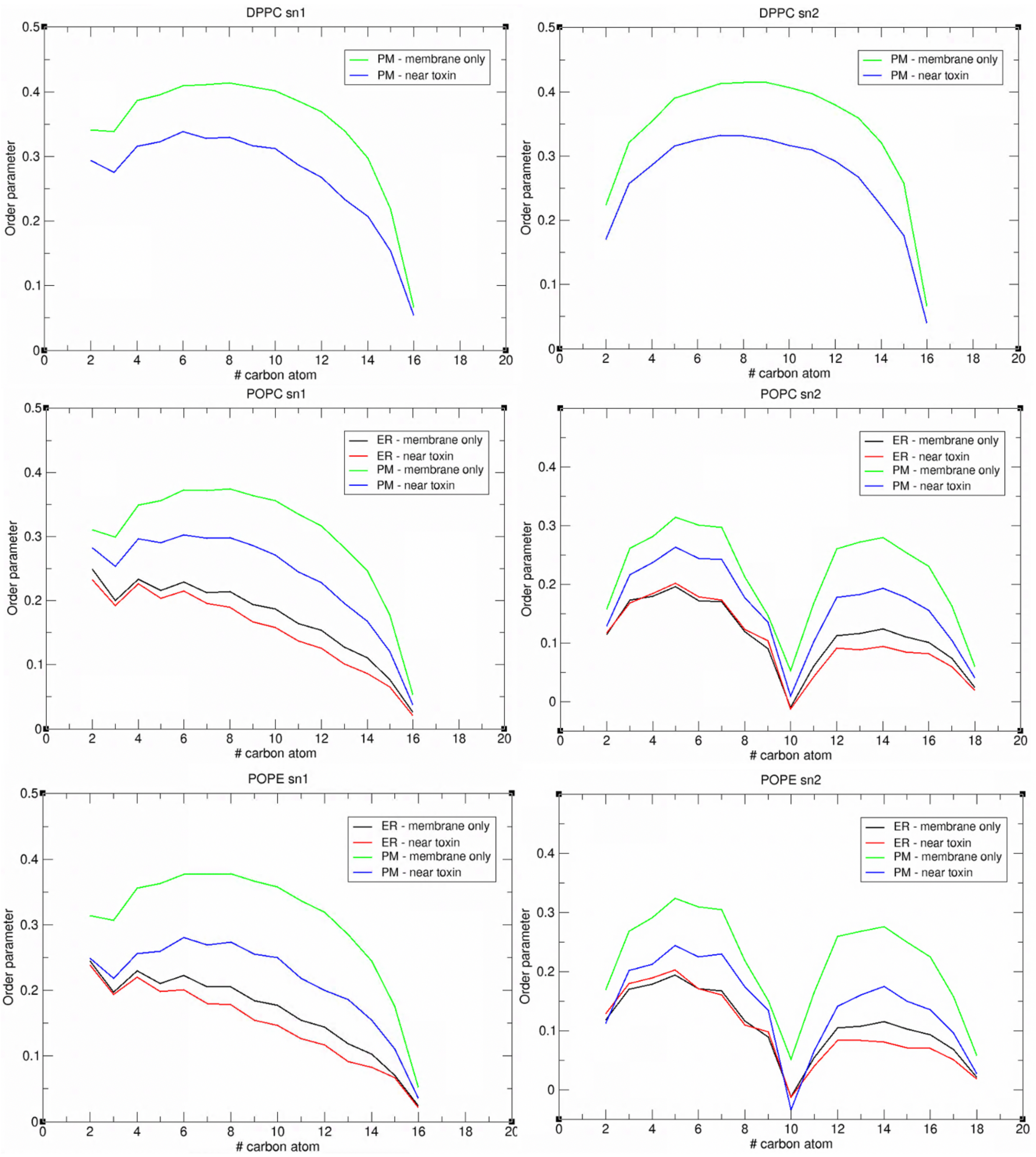


**Figure S4**. Tail order parameters of each phospholipid making up the ER and PM in unbiased simulations with the toxin and membrane only simulations.

### SUPPORTING MATERIAL

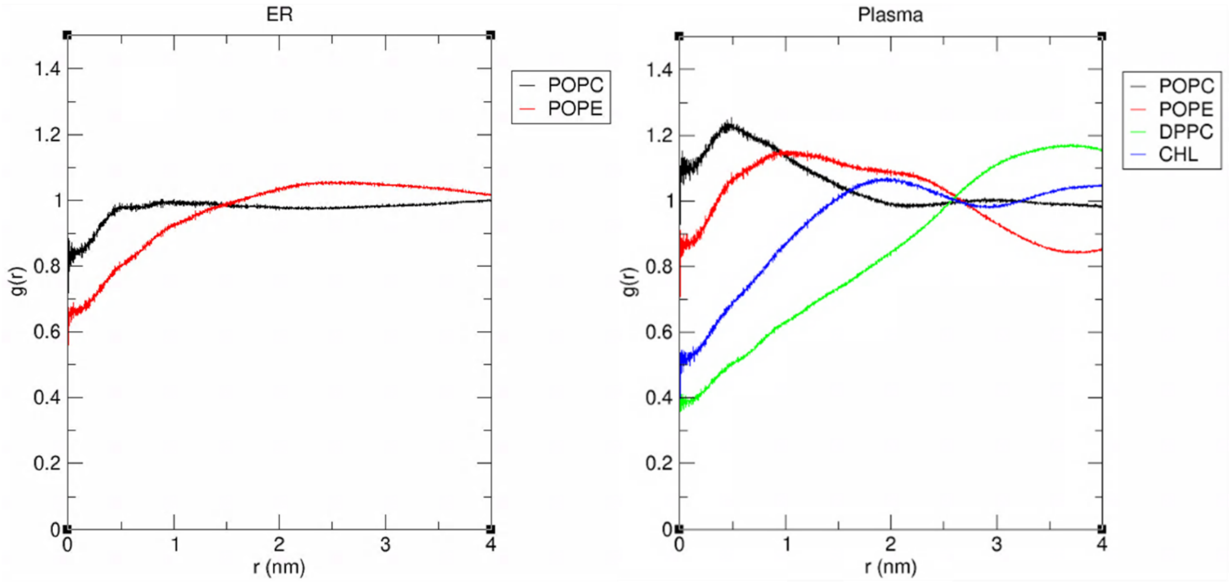


**Figure S5**. Radial distribution function of mycolactone with the center of mass of lipids making up the ER and PM in unbiased simulations.

**Table S1.** Average interaction potential energy of mycolactone with different components of the system.

| Mycolactone interaction energies (kcal mol$^{-1}$) | | | | |
|---|---|---|---|---|
| | Lipids | Water | Intramolecular | Total |
| Large ER | -103.68 ± 1.17 | -32.10 ± 0.84 | -10.69 ± 0.48 | -146.47 |
| Large PM | -108.28 ± 1.63 | -20.99 ± 0.91 | -11.49 ± 0.45 | -140.76 |

**Table S2.** Average interaction potential energy between lipids of the large ER membrane and large PM in systems with the toxin and systems without*.

| Lipid-Lipid interaction energy (kcal mol$^{-1}$) | | | | | |
|---|---|---|---|---|---|
| | Near the toxin | Far from the toxin | $\Delta_{(near-far)}$ | Membrane without toxin | $\Delta_{(near-without)}$ |
| ER | -144.33 ± 0.07 | -208.35 ± 0.29 | 64.02 | -196.80 ± 0.05 | 51.72 |
| PM | -90.82 ± 0.06 | -168.94 ± 0.14 | 78.12 | -143.16 ± 0.09 | 52.34 |

### SUPPORTING MATERIAL

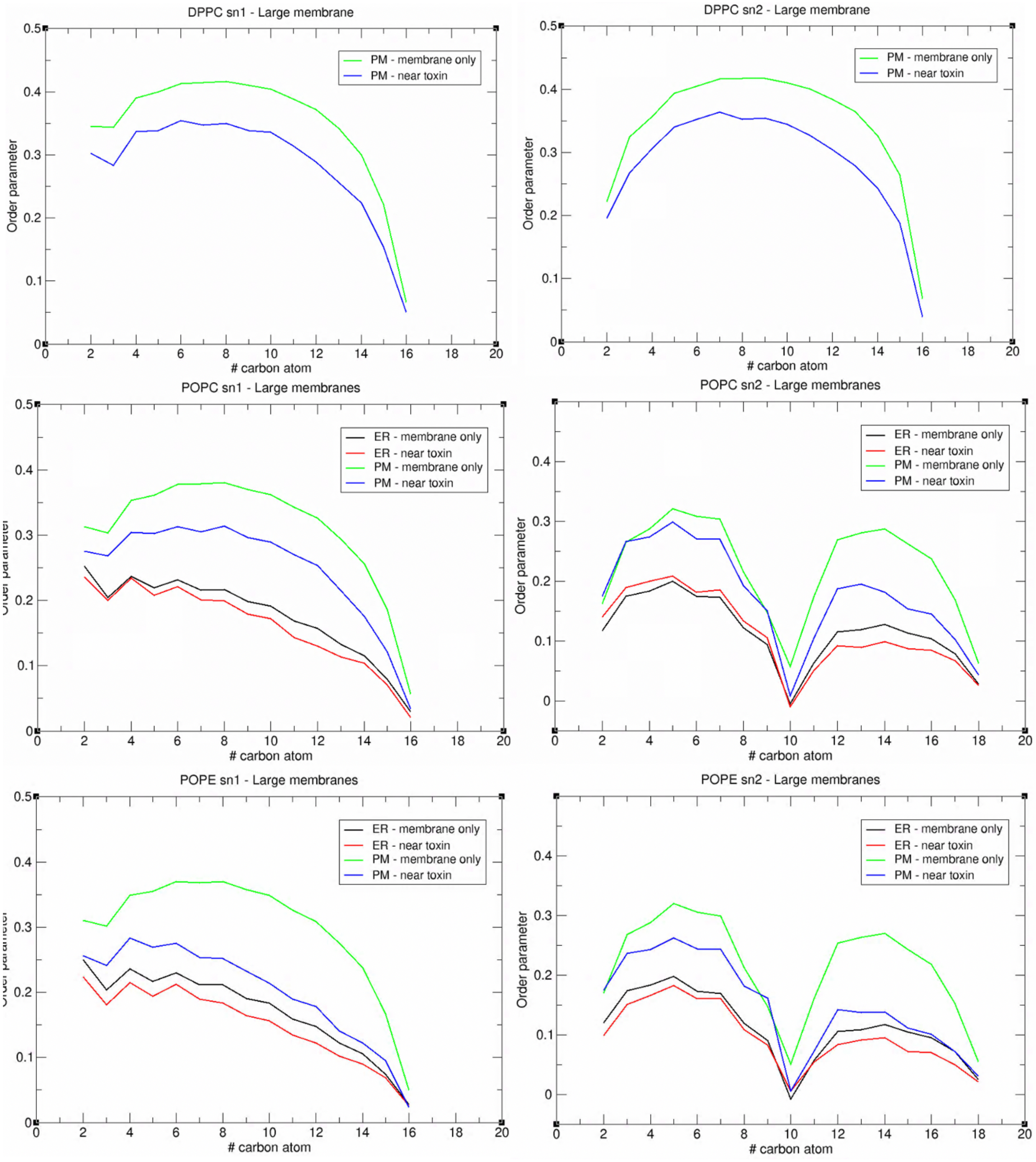


**Figure S6**. Tail order parameters of each phospholipid making up the large ER and large PM.

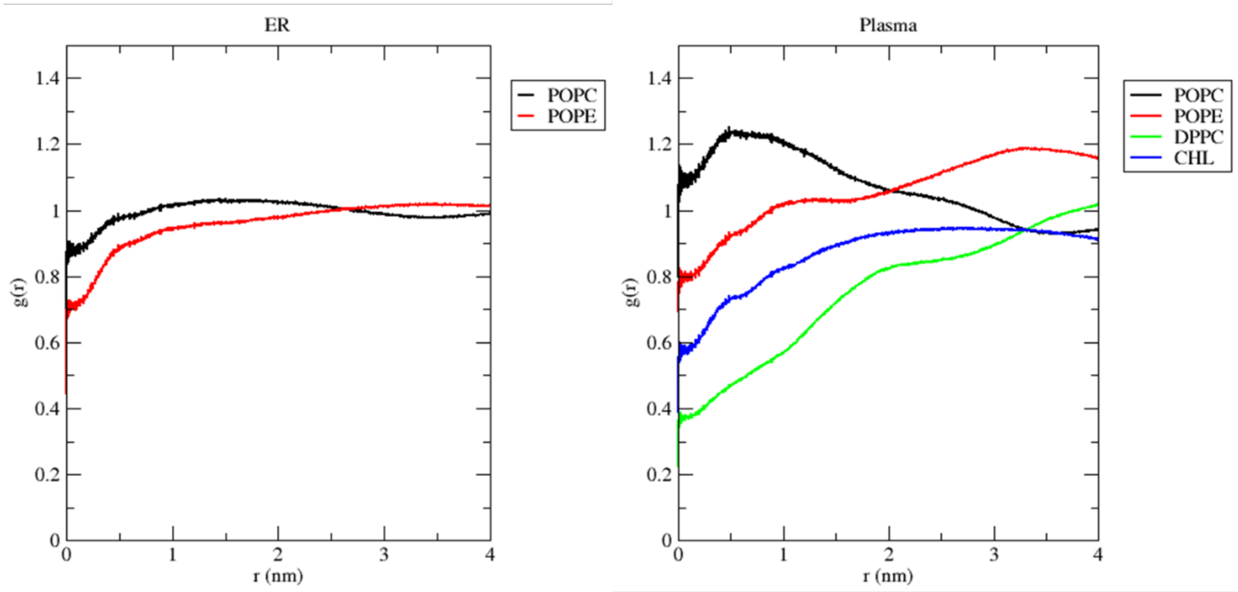


**Figure S7**. Radial distribution function of mycolactone with the center of mass of lipids making up the large ER and large PM in unbiased simulations.
